# A mannitol-based buffer improves single-cell RNA sequencing of high-salt marine cells

**DOI:** 10.1101/2023.04.26.538465

**Authors:** Tal Scully, Allon Klein

## Abstract

Single-cell RNA sequencing (scRNA-seq) enables discovery of novel cell states by transcriptomic profiling with minimal prior knowledge, making it useful for studying non-model organisms. For most marine organisms, however, cells are viable at a higher salinity than is compatible with scRNA-seq, impacting data quality and cell representation. We show that a low-salinity phosphate buffer supplemented with D-mannitol (PBS-M) enables higher-quality scRNA-seq of blood cells from the tunicate *Ciona robusta*. Using PBS-M reduces cell death and ambient mRNA, revealing cell states not otherwise detected. This simple protocol modification could enable or improve scRNA-seq for the majority of marine organisms.

## Background

Single-cell RNA sequencing (scRNA-seq) represents several methods for measuring the mRNA composition of individual cells at genome scale. These methods enable identification and characterization of previously unknown cell types with minimal prior knowledge. scRNA-seq has led to new discoveries in well-studied animals, but it is particularly useful when studying non-model organisms where the wide diversity of cell types is unexplored (1). At present, the highest-quality scRNA-seq methods require live cells, prepared in suspension after dissociation from tissues (2–5).

Salinity introduces a challenge for droplet-based scRNA-seq of marine organisms. In droplet-based methods, the buffer in which cells are loaded must be compatible with the first step in scRNA-seq, a reverse transcription (RT) of mRNA to make complementary DNA (cDNA). This reaction is carried out using variants of the Moloney Murine Leukemia Virus (MMLV) reverse transcriptase, which is efficient at the osmolarity of vertebrate cells (approximately 300 mOsm/L) (4,5). Most marine organisms — invertebrates, protists, and some vertebrates — are osmoconformers, with an internal osmolarity close to that of sea water (approx. 1000 mOsm/L) (6–10). RT enzyme efficiency drops below 10% at salt concentrations above 200 mM (400 mOsm/L) (4), so cells isolated from marine osmoconformers cannot be introduced into droplet-based scRNA-seq using native-salinity buffers.

Currently, a standard approach for scRNA-seq of marine cells is to dilute live cells to a lower salinity (≤300 mOsm/L) before rapid droplet-based encapsulation and subsequent RT. Datasets from multiple marine organisms have been collected in this way, including from tunicates, echinoderms, arthropods, cephalopods, and cnidarians (11–17). This strategy is premised on cells surviving the decreased osmotic pressure over the time required to complete encapsulation, but some cells might react to, or die from, the osmotic shock. Dying cells could be missing from the dataset and contribute ambient mRNA to other cells, and other cells may exhibit altered transcriptomic states.

We report on a simple method to prepare cells for droplet-based scRNA-seq that satisfies both the osmolarity requirements of marine cells and the salinity requirements of the RT. We benchmark the method with blood cells from the tunicate *Ciona robusta*, which we show are sensitive to osmotic shock. We expect that our approach will be broadly useful for single-cell study of marine organisms.

## Results and Discussion

This study was motivated by an interest in carrying out scRNA-seq on *C. robusta* blood. As a standard control prior to scRNA-seq, *C. robusta* blood cells were harvested and their viability was evaluated by Trypan blue staining after resuspension in PBS (∼330 mOsm/L) (18), a low-salt buffer routinely used with droplet-based scRNA-seq. We observed an 81% loss of live cells in PBS, as compared to a 40% loss after resuspension in buffered Ca^2+^- and Mg^2+^-free artificial seawater (CMF-ASW, ∼1000 mOsm/L). This loss mostly occurred within 1 minute of resuspension (**Fig 1a**).

**Fig. 1.**
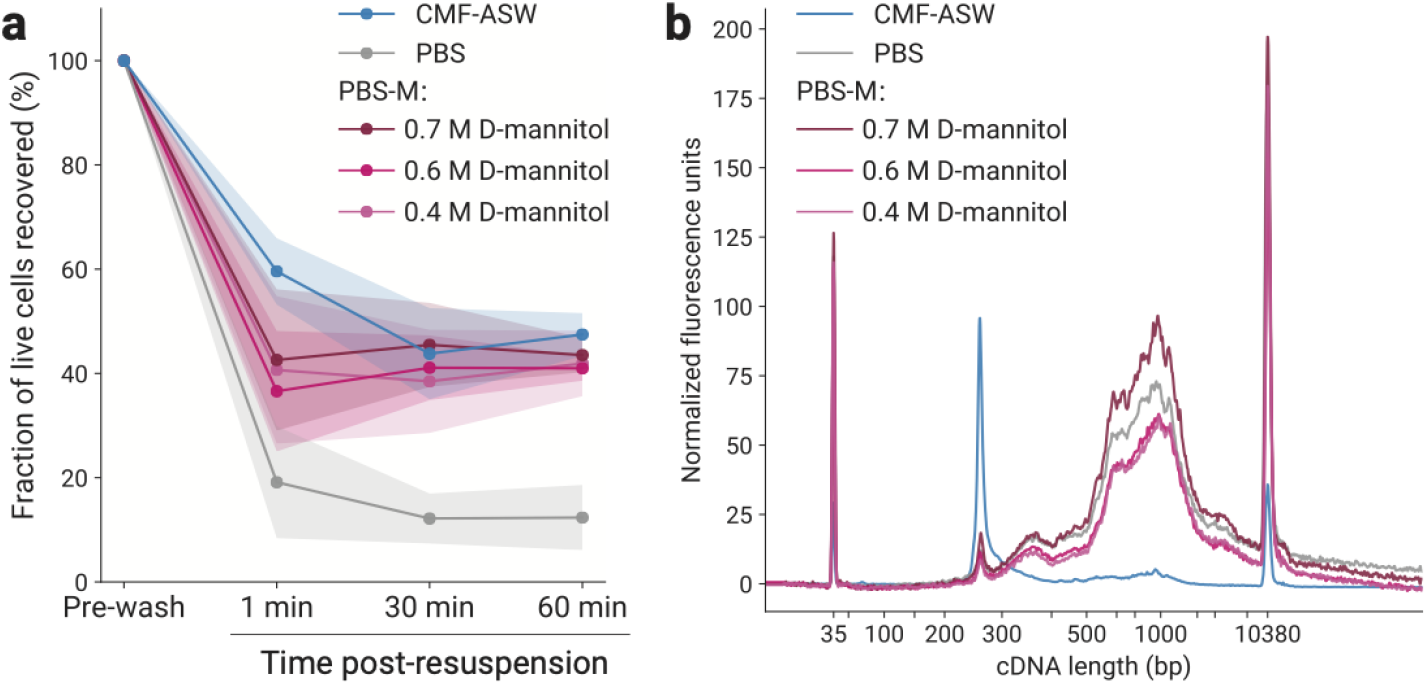
PBS-M buffer improves cell viability and does not inhibit reverse transcription. **a**, The fraction of live *C. robusta* blood cells recovered after resuspension in CMF-ASW, PBS, or PBS-M buffers. Data is averaged over 3–4 replicates. Shaded regions indicate 95% confidence intervals. **b**, Capillary electrophoresis traces of amplified cDNA from L1210 cells prepared in CMF-ASW, PBS, or PBS-M buffers. Peaks at 35 bp and 10380 bp are size markers.

We evaluated buffers for compatibility with scRNA-seq and for the ability to maintain cell viability. One such buffer consisted of 0.7 M D-mannitol dissolved in 1X PBS (PBS-M). D-mannitol is used as an osmotic agent in medical settings and several electroporation protocols (19–22). At 0.7 M concentration, it contributes 700 mOsm/L, thus PBS-M has comparable osmolarity to sea water (∼1000 mOsm total) with an ionic strength similar to PBS. When we resuspended *C. robusta* blood cells in PBS-M, there was no significant increase in cell loss as compared to resuspension in CMF-ASW (**Fig 1a**). This was consistent at reduced D-mannitol concentrations of 0.4 and 0.6 M.

PBS-M is compatible with the RT reaction required for scRNA-seq. To show this, we performed RT and subsequent PCR amplification on mouse leukemia L1210 cell lysates in either PBS, CMF-ASW, or PBS-M buffers with varying mannitol concentrations (0.4, 0.6, and 0.7 M). Capillary electrophoresis of the amplified RT products showed that reaction efficiency was poor in CMF-ASW, as expected due to high salinity (23). However, the quantity and size distribution of the resulting cDNA was similar between PBS and all tested concentrations of PBS-M (**Fig 1b**).

We next tested PBS-M for use in scRNA-seq. Blood cells from five adult animals were collected, pooled, and split equally into two samples that were prepared in tandem and jointly analyzed on a 10X Genomics Chromium scRNA-seq system (**Fig 2a**). The first sample was prepared by standard dilution as in (11): cells were pelleted and resuspended in CMF-ASW, then diluted >4.3-fold with water prior to droplet encapsulation. For the second sample, cells were instead resuspended in PBS-M, then diluted with PBS-M instead of water prior to droplet encapsulation. A target of 6,000 cells were collected for each aliquot. scRNA-seq libraries were sequenced and analyzed to evaluate evidence of cell death and cell state changes resulting from sample preparation.

**Fig. 2.**
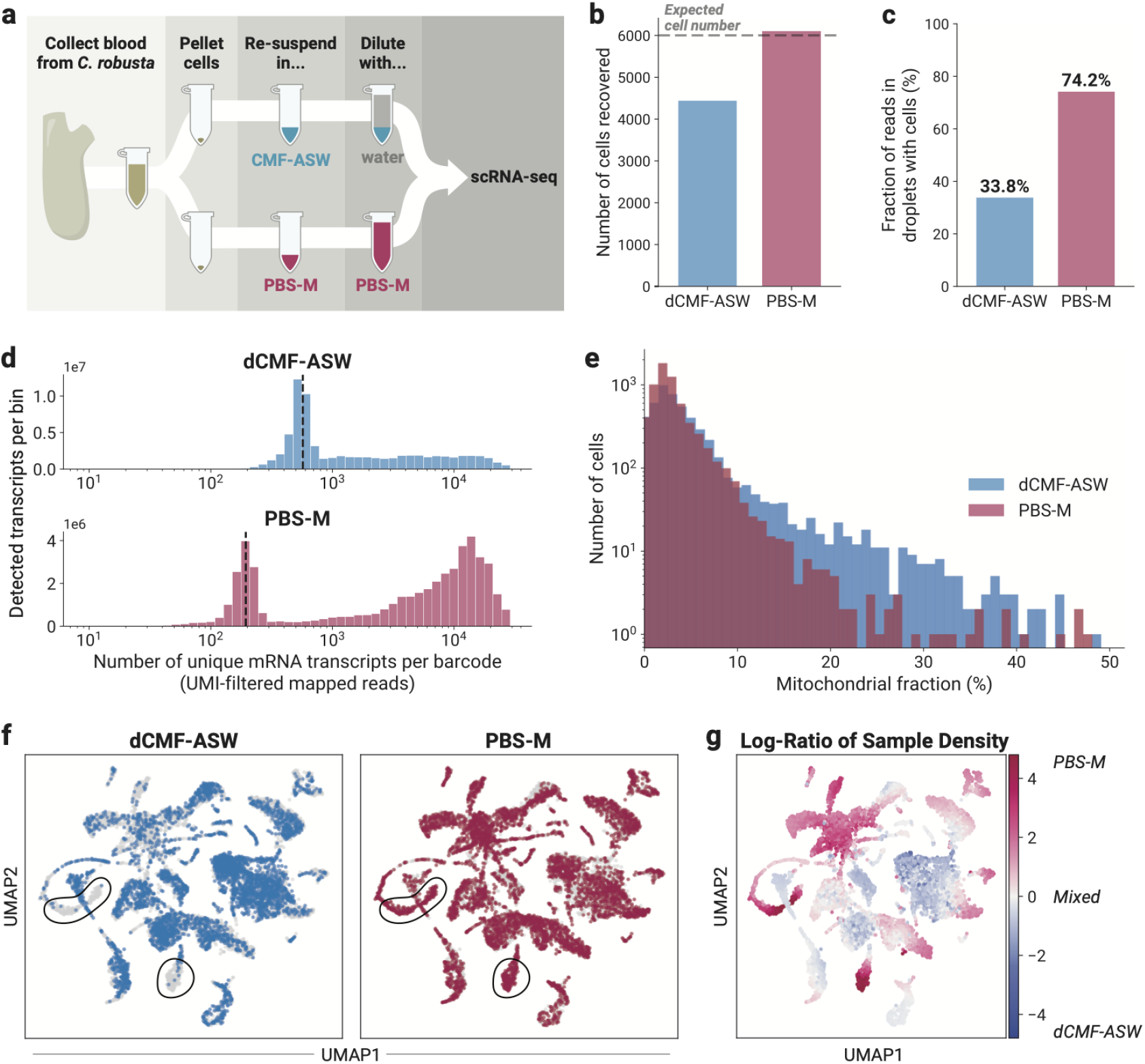
PBS-M buffer improves scRNA-seq data quality and recovers missing cell states in *C. robusta* blood. **a**, Experimental set-up for testing the effect of cell suspension buffer on scRNA-seq data quality. Blood from five animals was used. **b**,**c**, Preparing cells by standard dilution of high-salt buffer (dCMF-ASW) or PBS-M renders differences in: **b**, the number of single cell barcodes passing the scRNA-seq data filters, and **c**, the fraction of mapped reads whose barcodes pass the filters. Cells are filtered by the number of transcripts detected and mitochondrial fraction. **d**, The number of mRNA molecules detected per bin, with cell barcodes binned based on their transcript count. The dashed line shows the median number of transcripts in non-cell barcodes (571 transcripts/barcode in dCMF-ASW and 194 transcripts/barcode in PBS-M). These values quantify the background resulting from free-floating mRNA, which is reduced in PBS-M. **e**, Histogram of the mitochondrial fraction per cell, an indicator of cell lysis. **f**, Joint UMAP embedding of cells prepared in dCMF-ASW or PBS-M buffers, with cells colored by sample. Left: dCMF-ASW cells are marked in blue with the rest of the cells in grey. Right: PBS-M cells are marked in pink with the rest of the cells in grey. Cell states strongly depleted in the dCMF-ASW sample are circled in black. **g**, Joint UMAP embedding with cells colored by the log-ratio of the relative local sample density. The coloring represents the amount of local enrichment of cells from PBS-M (pink) or dCMF-ASW (blue) samples.

Several metrics indicate that PBS-M led to fewer dying cells than diluted CMF-ASW (dCMF-ASW). (i) Cell counts: we detected 6,108 viable cells in the PBS-M sample as compared to 4,443 for dCMF-ASW, even though we prepared the same number of cells for each condition (**Fig 2b**). (ii) Ambient mRNA: dying cells release their mRNA, leading to increased ambient mRNA levels that are detectable in empty droplets. The PBS-M sample had considerably lower levels of such ambient mRNA as compared to dCMF-ASW. This is seen from both the increased fraction of transcript counts from viable cells (**Fig 2c**) (74.2% for PBS-M vs. 33.8% for dCMF-ASW), and from the decreased median number of transcript counts in empty droplets (194 transcripts/droplet for PBS-M vs. 571 for dCMF-ASW) (**Fig 2d**). (iii) Mitochondrial transcripts: cells dying during scRNA-seq sample preparation lose nuclear-derived mRNA but retain mitochondrial-derived mRNA, leading to an increased fraction of mitochondrial transcript counts (*f*_*mito*_) (24). The PBS-M sample has 0.5% cells with *f*_*mito*_ >20%, compared to 3.6% for dCMF-ASW (**Fig 2e**). Overall, these metrics demonstrate that basic data quality metrics are greatly improved by using PBS-M.

In *C. robusta* blood, the cells prepared in PBS-M also reveal transcriptional states which are depleted or entirely absent when using dCMF-ASW. To visualize this, we generated a joint UMAP embedding and colored cells by preparation method (**Fig 2f**) and by a measure of local sample enrichment or depletion (the log-ratio of the number of nearest neighbors that belong to the PBS-M sample vs. the dCMF-ASW sample) (**Fig 2g**). These plots reveal cell clusters that were under-represented in the dCMF-ASW sample, potentially corresponding to cell types that are sensitive to osmotic shock.

## Conclusions

scRNA-seq has been revolutionary in enabling the discovery of new cell types, developmental and regenerative cell states, and complex immune responses. Even in well-studied mammalian systems, scRNA-seq has revealed unappreciated diversity of cell phenotypes (25,26). The number of cell states in mammals is dwarfed by the sheer number present across invertebrates. Each invertebrate species hosts a diversified repertoire of cell types, and few invertebrate animal tissues are defined at a cellular level. For instance, invertebrate immune cells are still largely described by cell morphology, with few or no molecular markers available to discriminate between cell types (27,28). Thus, scRNA-seq can short-cut years of work to identify and characterize cell types, enabling cell type comparisons across species.

Despite the promise of the technique, scRNA-seq was developed and optimized largely for vertebrates and is not compatible with high salinity buffers that mimic the body fluids of most marine animals (23). We have shown that using the low-salt, high-osmolarity PBS-M buffer for scRNA-seq of *C. robusta* blood cells greatly improves data quality by significantly reducing cell death. We saw two types of improvement: First, the baseline data quality obtained was considerably higher. Second, PBS-M let us observe certain cell states which were under-sampled or absent when using the standard dilution approach to scRNA-seq of marine cells.

With these advantages, some caveats are due. First, scRNA-seq may lose cell states independent of the buffers used. This is particularly true for large or fragile cells in tissues that are difficult to dissociate (29). Second, we cannot rule out the possibility that PBS-M may itself induce transcriptomic changes in some circumstances. In the tunicate *Botryllus schlosseri*, for example, D-mannitol and other sugars can inhibit blood cell phagocytic activity, which could over time lead to transcriptional changes in these cells (30).

We nonetheless expect that PBS-M should be suitable for preparing cells for scRNA-seq analysis from most, if not all marine invertebrate species and cell types. Certain species, such as some annelids, may require different D-mannitol concentrations to match their body osmolarity (10). To this end, we recommend direct assessment of cell viability in PBS-M prior to performing scRNA-seq (as in **Fig 1a**). Beyond this test, the use of PBS-M is simple. It could be broadly useful for generating high-quality scRNA-seq data from non-model marine organisms, better enabling researchers to explore the diversity of marine cell types.

## Methods

### Blood Cell Collection

Adult *Ciona robusta* supplied by M-REP (Carlsbad, California, United States of America) were kept at 17–18 °C in artificial seawater (ASW, Instant Ocean SS15-10) and fed daily with 15 mL of Phyto-Feast (Reef Nutrition). All animals were analyzed at 1–2 days after arriving in the lab.

Adults were relaxed by incubating for 3–16 hours at 4 °C in about 400 mL ASW with approximately 5 g of 99% L-menthol crystals (Thermo Scientific A10474) (3 adults per 400 mL volume). Relaxed adults were dissected to expose the endostyle and large blood vessel on the ventral side of the animal. Blood was drawn from this vessel using a zero dead space tuberculin syringe with a 25G needle, then transferred to a microcentrifuge tube on ice which had previously been coated with bovine serum albumin (BSA) by washing with 0.5 mL of 10% (v./v.) BSA (Sigma 9048-46-8) in Dulbecco’s phosphate-buffered saline (DPBS, Gibco 14200075).

To resuspend blood cells in alternate buffers, pooled blood was split into several tubes coated with 10% (v./v.) BSA in DPBS. Cells were pelleted by spinning at 800 g for 10 min at 4 °C in a swinging bucket centrifuge (Eppendorf Centrifuge 5810R). The supernatant was gently removed, and cells were resuspended in the appropriate buffer.

### Cell Viability Assay

Pooled blood cells from 9 adults were counted and their viability was assessed using an automatic cell counter (Bio-Rad TC20) with Trypan Blue staining. The cell suspension was evenly split and resuspended in one of 5 buffers: PBS, Ca^2+^- and Mg^2+^-free artificial seawater (CMF-ASW; 0.5 M NaCl, 9 mM KCl, 5 mM HEPES, pH 7.4), or PBS-M with varying mannitol concentrations (0.7 M, 0.6 M, or 0.4 M D-mannitol in PBS). Cell concentration and viability were measured with Trypan Blue at the time points post-resuspension indicated in **Fig 1a**.

In **Fig 1a**, the percent of live cells recovered was calculated by dividing the live (Trypan Blue negative) cell count at each time point by the live cell count before pelleting and resuspension. The data in **Fig 1a** includes a total of 3 replicates of CMF-ASW and 4 replicates of all other buffer conditions. Samples were collected across two separate days (1–2 replicates per condition per day).

### Reverse Transcription Efficiency Assay

L1210 cells (ATCC CCL-219) were grown in suspension in DMEM (Thermo Fisher 10566016) supplemented with 10% FBS and 1X Penicillin-Streptomycin (Thermo Fisher 15140122). The cells in culture media were pelleted in a swinging bucket centrifuge (Eppendorf Centrifuge 5810R) at 400 g for 4 min at 4 °C, then washed 3 times with ice-cold DPBS. The cells were pelleted and resuspended in one of 5 buffers: PBS, CMF-ASW, or PBS-M with varying mannitol concentrations (0.7 M, 0.6 M, or 0.4 M D-mannitol in PBS). Cells were diluted in the same buffer to a concentration of 1,000 cells/μL.

Each cell suspension was mixed 1:1 with a 2X reverse transcription (RT) mix, such that the final solution consisted of 500 cells/μL, 1X Maxima RT buffer (Thermo Scientific EP0751), 0.5 mM of each dNTP (Thermo Scientific R0192), 2.5 μM poly-T primer (5′-CTCACTATAGGGTGTCGGGTGCAGNNNNNNTTTTTTTTTTTTTTTTTTTV-3′), 25 μM template switching oligo (5′-AAGCAGTGGTATCAACGCAGAGTACATrGrGrG-3′), 1 U/μL RiboLock RNase inhibitor (Thermo Scientific EO0381), 0.3% IGEPAL NP-40 detergent (Sigma I8896-50ML), and 10 U/μL Maxima H minus RT enzyme (Thermo Scientific EP0751). The RT reaction was carried out at 50 °C for 60 min followed by inactivation at 85 °C for 5 min. RT samples were diluted by 1/4 in nuclease free water to reduce the salt concentration, then purified with a 1.2X volume of AMPure (Beckman Coulter, A63881) following the manufacturer’s instructions.

cDNA was amplified by PCR using KAPA HiFi HotStart ReadyMix (Roche Sequencing, KK2602) and 0.5 μM each of forward (5′-CTCACTATAGGGTGTCGGGTGCAG-3′) and reverse (5′-AAGCAGTGGTATCAACGCAGAGTACAT-3′) amplification primers. Amplified cDNA was purified once more using a 1.2X volume of AMPure. Finally, cDNA was diluted to approximately 3 ng/μL and run on the 2100 Agilent Bioanalyzer using the High Sensitivity DNA Kit (Agilent 5067-4626). The measured fluorescence units in **Fig 1b** have been corrected for the dilution factor.

### Single-cell Barcoding, Library Preparation, and Sequencing

Pooled blood cells from 5 adults were collected as described above, split into two aliquots, and then resuspended in either CMF-ASW or PBS-M (0.7 M D-mannitol in PBS) at a concentration of 1,000–2,000 cells/μL. Cell encapsulation, RT, cDNA amplification, and library preparation was done using the Chromium Next GEM Single Cell 3′ v3.1 Kits (10X Genomics 1000123, 1000127, and 1000190), following the manufacturer’s guidelines except for one step. When preparing the RT mix prior to cell encapsulation, the manufacturer protocol instructs to add nuclease-free water, then to add the cell suspension. For the cells in CMF-ASW, we followed these directions (diluted CMF-ASW, or dCMF-ASW); for the PBS-M sample, we added PBS-M instead of adding nuclease-free water.

Libraries were sequenced with Illumina NovaSeq 6000 SP, and a cell by gene count matrix was generated from FASTQ files using 10X Genomics Cell Ranger 6.1.2. For a reference genome, the HT2019 assembly with KY21 gene models from the Ghost Database (31,32) was used in combination with mitochondrial genes from the Ensembl KH genome assembly (33) (gene IDs listed in Additional File 1). The available GFF3 file of the KY21 gene models was reformatted as a GTF file using GffRead (34) in order to be compatible with 10X Cell Ranger.

### Cell Filtering and Data Quality Measures

Viable cells were filtered from background based on (i) the number of unique mRNA molecules (UMI-filtered mapped reads, or UMIs) detected, (ii) mitochondrial fraction, and (iii) scoring low for likelihood of being a doublet. For (i), a UMI count above a threshold was required. These thresholds were set based on a clear separation from background as seen by visual inspection of the plots in **Fig 1d**. The threshold values used were 2000 UMIs for dCMF-ASW and 800 for PBS-M. For (ii), a mitochondrial fraction (*f*_*mito*_) below 10% was required for both samples. For (iii), likely doublets were identified using the SCRUBLET package (35). All cell barcodes are included in **Fig 2d**, and only cell barcodes passing the UMI threshold are plotted in **Fig 1e**.

The median UMIs/barcode in empty droplets for each sample, as marked with a dashed line in **Fig 2d**, was calculated across empty droplets. Empty droplets were defined as barcodes which did not pass the UMI threshold (<800 UMIs for PBS-M, <2000 UMIs for dCMF-ASW) and which had ≥10 UMIs/barcode.

### Dimensionality Reduction

Preprocessing and dimensionality reduction were done using Scanpy (36) (version 1.8.1) using default parameters except where noted below. Genes expressed in fewer than 5 cells were excluded. Counts were normalized (pp.normalize_total) and log transformed (pp.log1p), and highly variable genes were identified (pp.highly_variable_genes) as described in (37). Principle component (PC) analysis was performed (tl.pca, use_highly_variable=True) on z-scored data (pp.scale). A nearest neighbor network (pp.neighbors, n_neighbors=10) was generated based on Euclidean distance in PC space using the first 50 PCs. This network was used for UMAP visualization (tl.umap) (38).

### Log-Ratio of Sample Density

The log-ratio of sample density is a measure of enrichment or depletion of one sample relative to another in gene expression space. We adapted the measure defined in (25) as follows: the 200 nearest neighbors of each cell *i* were identified in PC space using the first 50 PCs (Euclidean distance). A cell’s “neighborhood” includes itself and its *k* nearest neighbors. Let *N*_*i,dCMF*−*ASW*_ (*k*) and *N*_*i,PBS*−*M*_ (*k*) be the number of cells within the neighborhood of cell *i* from the dCMF-ASW and the PBS-M samples, respectively. We used *k*=200 and drop the argument *k* below. For each cell *i*, a numerical representation of the log-scaled relative sample density, shown in **Fig 2g**, was calculated as

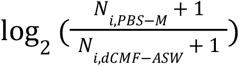

These values were plotted in a UMAP generated by Scanpy (pl.umap, sort_order=False).

## Declarations

### Ethics approval and consent to participate

Not applicable

### Consent for publication

Not applicable

### Availability of data and materials

The datasets used and/or analyzed during the current study are available from the corresponding author on reasonable request.

### Authors’ contributions

TS carried out the experiments and data analysis. AM supervised the work. TS and AM wrote the manuscript. Both authors read and approved the final manuscript.

## Acknowledgments

The authors would like to thank Andrew Murphy for building aquariums and supporting the animals; Ignas Mazelis for assisting with the RT assay; and Sean McGeary for comments on this manuscript. We also thank the Single Cell Core at Harvard Medical School for assisting with the scRNA-seq sample preparation and providing 10X reagents, and the Bauer Core Facility at Harvard University for sequencing.

## Additional File 1

### Ensemble gene ID for mitochondrial genes

ENSCING00000025191

ENSCING00000025192

ENSCING00000025193

ENSCING00000025194

ENSCING00000025195

ENSCING00000025196

ENSCING00000025197

ENSCING00000025198

ENSCING00000025199

ENSCING00000025200

ENSCING00000025201

ENSCING00000025202

ENSCING00000025203

ENSCING00000025204

ENSCING00000025205

ENSCING00000025206

ENSCING00000025207

ENSCING00000025208

ENSCING00000025209

ENSCING00000025210

ENSCING00000025211

ENSCING00000025212

ENSCING00000025213

ENSCING00000025214

ENSCING00000025215

ENSCING00000025216

ENSCING00000025217

ENSCING00000025218

ENSCING00000025219

ENSCING00000025220

ENSCING00000025221

ENSCING00000025222

ENSCING00000025223

ENSCING00000025224

ENSCING00000025225

ENSCING00000025226

ENSCING00000025227

ENSCING00000025228

ENSCING00000025229

## References

1. Alfieri JM, Wang G, Jonika MM, Gill CA, Blackmon H, Athrey GN. A Primer for Single-Cell Sequencing in Non-Model Organisms. Genes [Internet]. 2022 Feb 19;13(2). Available from: http://dx.doi.org/10.3390/genes13020380

2. Wang X, Yu L, Wu AR. The effect of methanol fixation on single-cell RNA sequencing data. BMC Genomics. 2021 Jun 5;22(1):420.

3. Chen J, Cheung F, Shi R, Zhou H, Lu W, CHI Consortium. PBMC fixation and processing for Chromium single-cell RNA sequencing. J Transl Med. 2018 Jul 17;16(1):198.

4. Pan Y, Cao W, Mu Y, Zhu Q. Microfluidics Facilitates the Development of Single-Cell RNA Sequencing. Biosensors [Internet]. 2022 Jun 24;12(7). Available from: http://dx.doi.org/10.3390/bios12070450

5. Rosenberg AB, Roco CM, Muscat RA, Kuchina A, Sample P, Yao Z, et al. Single-cell profiling of the developing mouse brain and spinal cord with split-pool barcoding. Science. 2018 Apr 13;360(6385):176–82.

6. Bradley TJ. Osmoconformers. 2008 Dec 25 [cited 2023 Mar 21]; Available from: http://dx.doi.org/10.1093/acprof:oso/9780198569961.003.0005

7. Yancey PH. Nitrogen compounds as osmolytes. In: Fish Physiology. Academic Press; 2001. p. 309–41.

8. Dakin WJ. The Composition of the Blood of Aquatic Animals and its Bearings upon the Possible Conditions of Origin of the Vertebrates. Nature. 1931 Jul;128(3219):66–7.

9. Robertson JD. Osmotic constituents of the body-wall muscles of the hemichordate Balanoglossus clavigerus, the tunicate Ciona intestinalis, and the cephalochordate Branchiostoma lanceolatum. Comp Biochem Physiol A Physiol. 1987 Jan 1;87(2):363–73.

10. Generlich O, Giere O. Osmoregulation in two aquatic oligochaetes from habitats with different salinity and comparison to other annelids. Hydrobiologia. 1996 Oct 1;334(1):251–61.

11. Cao C, Lemaire LA, Wang W, Yoon PH, Choi YA, Parsons LR, et al. Comprehensive single-cell transcriptome lineages of a proto-vertebrate. Nature. 2019 Jul 18;571(7765):349–54.

12. Zhang T, Xu Y, Imai K, Fei T, Wang G, Dong B, et al. A single-cell analysis of the molecular lineage of chordate embryogenesis. Sci Adv [Internet]. 2020 Nov;6(45). Available from: http://dx.doi.org/10.1126/sciadv.abc4773

13. Paganos P, Voronov D, Musser JM, Arendt D, Arnone MI. Single-cell RNA sequencing of the Strongylocentrotus purpuratus larva reveals the blueprint of major cell types and nervous system of a non-chordate deuterostome. Elife [Internet]. 2021 Nov 25;10. Available from: http://dx.doi.org/10.7554/eLife.70416

14. Koiwai K, Koyama T, Tsuda S, Toyoda A, Kikuchi K, Suzuki H, et al. Single-cell RNA-seq analysis reveals penaeid shrimp hemocyte subpopulations and cell differentiation process. Elife [Internet]. 2021 Jun 16;10. Available from: http://dx.doi.org/10.7554/eLife.66954

15. Songco-Casey JO, Coffing GC, Piscopo DM, Pungor JR, Kern AD, Miller AC, et al. Cell types and molecular architecture of the Octopus bimaculoides visual system. Curr Biol [Internet]. 2022 Oct 26; Available from: http://dx.doi.org/10.1016/j.cub.2022.10.015

16. Chari T, Weissbourd B, Gehring J, Ferraioli A, Leclère L, Herl M, et al. Whole-animal multiplexed single-cell RNA-seq reveals transcriptional shifts across Clytia medusa cell types. Sci Adv. 2021 Nov 26;7(48):eabh1683.

17. Hu M, Zheng X, Fan CM, Zheng Y. Lineage dynamics of the endosymbiotic cell type in the soft coral Xenia. Nature. 2020 Jun;582(7813):534–8.

18. Gallagher SR. Recipes for commonly encountered reagents. Curr Protoc Essent Lab Tech. 2018 Nov;17(1):e24.

19. Shawkat H, Westwood MM, Mortimer A. Mannitol: a review of its clinical uses. Contin Educ Anaesth Crit Care Pain. 2012 Apr 1;12(2):82–5.

20. Christiaen L, Wagner E, Shi W, Levine M. Electroporation of transgenic DNAs in the sea squirt Ciona. Cold Spring Harb Protoc. 2009 Dec;2009(12):db.prot5345.

21. Takao D, Kamimura S. Single-cell electroporation of fluorescent probes into sea urchin sperm cells and subsequent FRAP analysis. Zoolog Sci. 2010 Mar;27(3):279–84.

22. Moisescu MG, Radu M, Kovacs E, Mir LM, Savopol T. Changes of cell electrical parameters induced by electroporation. A dielectrophoresis study. Biochim Biophys Acta. 2013 Feb;1828(2):365–72.

23. Gerard GF, D’Alessio JM. Reverse Transcriptase (EC 2.7.7.49). In: Burrell MM, editor. Enzymes of Molecular Biology. Totowa, NJ: Humana Press; 1993. p. 73–93.

24. Ilicic T, Kim JK, Kolodziejczyk AA, Bagger FO, McCarthy DJ, Marioni JC, et al. Classification of low quality cells from single-cell RNA-seq data. Genome Biol. 2016 Feb 17;17:29.

25. Zilionis R, Engblom C, Pfirschke C, Savova V, Zemmour D, Saatcioglu HD, et al. Single-Cell Transcriptomics of Human and Mouse Lung Cancers Reveals Conserved Myeloid Populations across Individuals and Species. Immunity. 2019 May 21;50(5):1317–34.e10.

26. Villani AC, Satija R, Reynolds G, Sarkizova S, Shekhar K, Fletcher J, et al. Single-cell RNA-seq reveals new types of human blood dendritic cells, monocytes, and progenitors. Science [Internet]. 2017 Apr 21;356(6335). Available from: http://dx.doi.org/10.1126/science.aah4573

27. Hartenstein V. Blood Cells and Blood Cell Development in the Animal Kingdom. Annu Rev Cell Dev Biol. 2006 Nov 9;22(1):677–712.

28. Longo V, Parrinello D, Longo A, Parisi MG, Parrinello N, Colombo P, et al. The conservation and diversity of ascidian cells and molecules involved in the inflammatory reaction: The Ciona robusta model. Fish Shellfish Immunol. 2021 Oct 20;119:384–96.

29. Briggs JA, Weinreb C, Wagner DE, Megason S, Peshkin L, Kirschner MW, et al. The dynamics of gene expression in vertebrate embryogenesis at single-cell resolution. Science [Internet]. 2018 Jun 1;360(6392). Available from: https://science.sciencemag.org/content/360/6392/eaar5780

30. Ballarin L, Cima F, Sabbadin A. Phagocytosis in the colonial ascidian Botryllus schlosseri. Dev Comp Immunol. 1994 Nov 1;18(6):467–81.

31. Satou Y, Tokuoka M, Oda-Ishii I, Tokuhiro S, Ishida T, Liu B, et al. A Manually Curated Gene Model Set for an Ascidian, Ciona robusta (Ciona intestinalis Type A). jzoo [Internet]. 2022 Mar [cited 2022 Mar 1];39(3). Available from: https://bioone.org/journals/zoological-science/volume-39/issue-3/zs210102/A-Manually-Curated-Gene-Model-Set-for-an-Ascidian-Ciona/10.2108/zs210102.short

32. Satou Y, Kawashima T, Shoguchi E, Nakayama A, Satoh N. An integrated database of the ascidian, Ciona intestinalis: towards functional genomics. Zoolog Sci. 2005 Aug;22(8):837–43.

33. Cunningham F, Allen JE, Allen J, Alvarez-Jarreta J, Amode MR, Armean IM, et al. Ensembl 2022. Nucleic Acids Res. 2022 Jan 7;50(D1):D988–95.

34. Pertea G, Pertea M. GFF Utilities: GffRead and GffCompare. F1000Res [Internet]. 2020 Apr 28;9. Available from: http://dx.doi.org/10.12688/f1000research.23297.2

35. Wolock SL, Lopez R, Klein AM. Scrublet: Computational Identification of Cell Doublets in Single-Cell Transcriptomic Data. Cell Syst. 2019 Apr 24;8(4):281–91.e9.

36. Wolf FA, Angerer P, Theis FJ. SCANPY: large-scale single-cell gene expression data analysis. Genome Biol. 2018 Feb 6;19(1):15.

37. Satija R, Farrell JA, Gennert D, Schier AF, Regev A. Spatial reconstruction of single-cell gene expression data. Nat Biotechnol. 2015 May;33(5):495–502.

38. McInnes L, Healy J, Melville J. UMAP: Uniform Manifold Approximation and Projection for Dimension Reduction [Internet]. arXiv [stat.ML]. 2018. Available from: http://arxiv.org/abs/1802.03426

